# Utp14 interaction with the Small Subunit Processome

**DOI:** 10.1101/280859

**Authors:** Joshua J. Black, Zhaohui Wang, Lisa M. Goering, Arlen W. Johnson

## Abstract

The SSU Processome (sometimes referred to as 90S) is an early stabile intermediate in the small ribosomal subunit biogenesis pathway of eukaryotes. Progression of the SSU Processome to a pre-40S particle requires a large-scale compaction of the RNA and release of many biogenesis factors. The U3 snoRNA is a primary component of the SSU Processome and hybridizes to the rRNA at multiple locations to organize the structure of the SSU Processome. Thus, release of U3 is prerequisite for the transition to pre-40S. Our lab proposed that the RNA helicase Dhr1 plays a crucial role in the transition by unwinding U3 and that this activity is controlled by the SSU Processome protein Utp14. How Utp14 times the activation of Dhr1 is an open question. Despite being highly conserved, Utp14 contains no recognizable domains, and how Utp14 interacts with the SSU Processome is not well characterized. Here, we used UV crosslinking and analysis of cDNA and yeast two-hybrid interaction to characterize how Utp14 interacts with the pre-ribosome. Moreover, proteomic analysis of SSU particles lacking Utp14 revealed that Utp14 is needed for efficient recruitment of the RNA exosome. Our analysis positions Utp14 to be uniquely poised to communicate the status of assembly of the SSU Processome to Dhr1 and possibly the exosome as well.

## INTRODUCTION

Ribosomes are the complex and dynamic molecular machines that decode genetic information into protein. In *Saccharomyces cerevisiae*, the ribosomal large subunit (LSU or 60S) is composed of three ribosomal RNA (rRNA) molecules (25S, 5.8S, 5S) and 46 ribosomal proteins (r-proteins), and the small subunit (SSU or 40S) consists of the 18S rRNA and 33 r-proteins (Ben-Shem et al. 2011). Ribosome synthesis begins in the nucleolus with co-transcriptional recruitment of assembly factors to the polycistronic 35S transcript. The 35S rRNA undergoes extensive modification and processing, coordinated with RNA folding and protein assembly, to generate the pre-40S and pre-60S particles, which are subsequently exported to the cytoplasm where the final steps of maturation occur (for recent reviews see (Kressler et al. 2017; Peña et al. 2017; Sloan et al. 2016)).

An early stabile intermediate of 40S assembly is the SSU Processome, a large complex of ∼6MDa containing the 5’-portion of the 35S rRNA transcript, the 5’-external transcribed spacer (5’-ETS), 18S and a portion of the internal transcribed spacer 1 (ITS1) (Kressler et al. 2017). The SSU processome also contains the U3 snoRNA and approximately 70 assembly factors (Chaker-Margot et al. 2015; Zhang et al. 2016). Although the SSU processome is sometimes referred to as the 90S pre-ribosomal complex, we will use the term SSU processome to avoid confusion with related particles that contain intact 35S rRNA. Recent high-resolution cryo-electron microscopy reconstructions of the SSU Processome from *S. cerevisiae* and the thermophilic fungus *Chaetomium thermophilum* reveal a highly splayed-open structure of the rRNA (Kornprobst et al. 2016; Chaker-Margot et al. 2017; Sun et al. 2017; Barandun et al. 2017; Cheng et al. 2017). The SSU Processome may represent a metastable intermediate of assembly, as particles with similar structure and composition have been purified from cells under various conditions including stationary phase in which ribosome biogenesis is largely repressed (Chaker-Margot et al. 2017; Barandun et al. 2017).

Progression of the SSU Processome to the pre-40S particle requires endonucleolytic cleavages at sites A_0_ and A_1_ within the 5’-ETS to generate the mature 5’-end of 18S and cleavage at site A_2_ within ITS1 (Kressler et al. 2017). This transition results in the release of most SSU Processome factors, and concomitant large-scale rearrangements of the RNA as the splayed open structure collapses into the more compact structure of the small subunit (Johnson et al. 2017; Heuer et al. 2017). What triggers the transition of the SSU Processome to a pre-40S is not yet known.

A primary feature of the SSU Processome is the U3 snoRNA (*SNR17A*/*B*) which hybridizes to multiple regions of the 5’-ETS as well as 18S rRNA to provide a scaffold for the initial folding of the pre-ribosomal RNA and assembly of the domains of the small subunit (Dragon et al. 2002; Sun et al. 2017; Barandun et al. 2017; Cheng et al. 2017). Importantly, U3 hybridizes to residues in the 5’-end of 18S that are involved in intramolecular base-pairing required to form the central pseudoknot, a critical RNA element that coordinates all domains of the small subunit (Henras et al. 2008). Consequently, U3 must be released to allow assembly of the central pseudoknot, and it is likely that the release of U3 is a principal driver of the RNA rearrangements that promote the transition from the SSU processome to the pre-40S particle. We previously provided evidence that the release of U3 is driven by the DEAH-box RNA helicase Dhr1 (Ecm16) (Sardana et al. 2015) whose stabile association and subsequent activation depends upon direct interactions with the SSU processome factor Utp14 (Zhu et al. 2016). However, how the timing of Dhr1 activation by Utp14 is controlled is not known. Utp14 joins the SSU Processome at a late stage of assembly, after the majority of the 3’-minor domain has been transcribed (Chaker-Margot et al. 2015; Zhang et al. 2016). Unlike the majority of SSU Processome factors, Utp14 remains associated with 20S rRNA (Sardana et al. 2013) suggesting it remains on the pre-ribosome during the transition from SSU Processome to pre-40S particle, however it is not present on cytoplasmic particles (Johnson et al. 2017; Heuer et al. 2017). Utp14 is a highly-conserved protein found throughout eukaryotes but contains no recognizable domains, and its interaction with the pre-ribosome has only recently begun to be revealed (Sardana et al. 2013; Zhu et al. 2016; Barandun et al. 2017; Cheng et al. 2017).

We sought to further characterize the interaction of Utp14 with the pre-ribosome to understand how it regulates Dhr1 activity in the context of the SSU Processome. Here, we used UV Crosslinking and Analysis of cDNA (CRAC) to identify the RNA binding sites of Utp14 and yeast 2-hybrid analysis to map domain interactions with assembly factors and small subunit r-proteins. In addition, we examined the protein and RNA composition of particles arrested with several Utp14 mutants. Our work is consistent with and extends recent structural and genetic analyses of the SSU Processome.

## RESULTS

### Utp14 binds multiple RNA elements in the SSU Processome

To determine the RNA binding sites of Utp14 we used a modified UV <u>cr</u>oss-linking and <u>a</u>nalysis of <u>c</u>DNA (CRAC) protocol (Granneman et al. 2009). UV irradiation induces covalent cross-links between amino acids and neighboring nucleic acids allowing for nucleotide-resolution of RNA binding sites of proteins. The C-terminal His6-tobacco etch virus (TEV) protease recognition site-protein A (HTP) tag was integrated into the genomic locus of *UTP14*. The HTP tag had no apparent effect on growth (data not shown). Cells were subjected to UV irradiation and RNAs crosslinked to Utp14-HTP were first affinity-purified via the protein A tag under native conditions followed by RNase treatment and a second step purification via its His6 tag under denaturing conditions. To verify that RNAs were crosslinked to Utp14, co-purifying RNAs were radio-labeled with ^32^P, separated by SDS-PAGE and autoradiographed (Figure 1A). The HTP-tagged sample contained a high molecular weight radiolabeled band that was not present in the untagged control. This species was excised, crosslinked RNAs were released from Utp14 by proteinase K digestion, and the crosslinked RNAs were sequenced following library preparation (see Materials and Methods). Whereas the CRAC protocol involves ligation of oligonucleotides to both ends of the RNA followed by reverse transcription and amplification, we ligated a single oligonucleotide to the 3’-ends of the RNAs, followed by reverse transcription, circularization of the resulting product and amplification. This strategy results in a characteristic drop-off of the reverse transcriptase on the 5’-end of the cDNA product where an amino acid was crosslinked to the RNA substrate.

**Figure 1.**
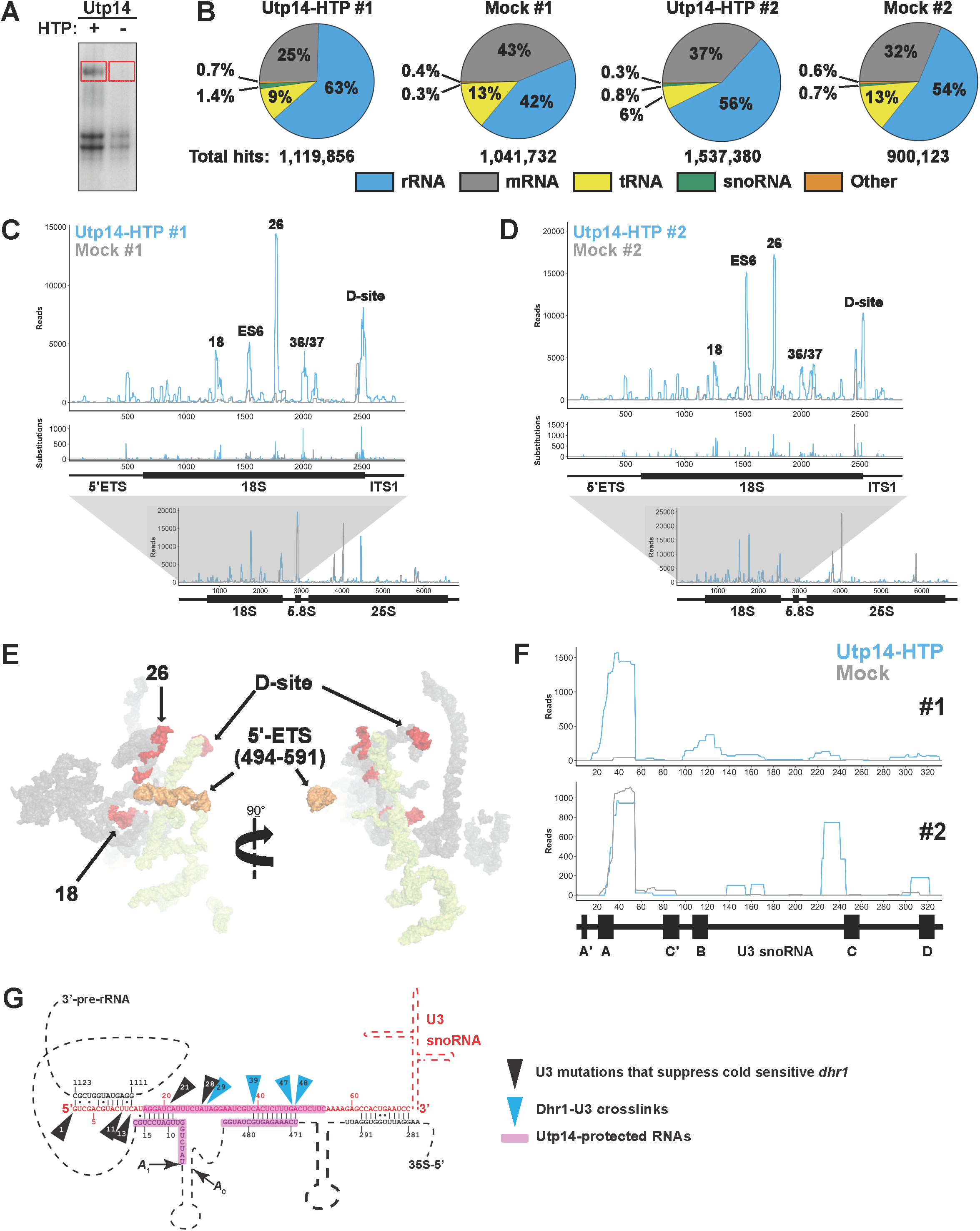
Utp14 crosslinks to multiple regions within the pre-18S rRNA. (A) A representative autoradiograph of ^32^P-labelled RNAs crosslinked to Utp14-HTP (+, AJY4051) and mock (-, BY4741). Red boxes indicate the regions of the membrane that were excised and used in library preparation. (B) Percentages of RNA composition grouped by class are shown for both CRAC replicates. Total aligned reads corresponding to each sample are shown below. (C, D) The number of reads (top) and substitutions (middle) are shown against nucleotide position within the pre-18S rRNA (*RDN18-1*). The number of reads against nucleotide position within the 35S rRNA (*RDN37-1*) are shown below. Utp14-HTP is shown in light blue, and the mock is shown as grey. Two independent biological replicates are shown. (E) Utp14 crosslinks within 18S rRNA (red) and 5’-ETS (orange) mapped to a recent structure of the SSU Processome. RNAs that were not crosslinked with Utp14 are shown in surface representation for18S rRNA (grey) and 5’-ETS (yellow) (PDB: 5WYJ). (F) The number of reads mapping to positions in U3 (*snR17A*) for datasets #1 (top) and #2 (bottom). A cartoon of U3 is shown below the plots. (G) A cartoon of U3 hybridization to the rRNA within the SSU Processome displaying Dhr1 crosslinks (blue triangles) and mutations of U3 that suppress a cold-sensitive Dhr1 mutant (black triangles) (Sardana et al. 2015) in relation to Utp14 crosslinks to U3 and the rRNA (magenta highlights). Relevant processed data are reported in Supplemental File 1.

Utp14-HTP crosslinked RNAs were enriched for rRNA and snoRNAs compared to the mock in both replicates (Figure 1B). Despite the lower level of rRNA enrichment in the second replicate, both datasets showed specific hits within rRNA, mapping primarily to pre-18S rRNA within 35S rRNA (Figure 1C and 1D; bottom). Utp14 crosslinked to multiple RNA elements within the pre-18S rRNA (Figures 1C and 1D; top). The highest read densities corresponded to nucleotides spanning helix 26es7 (hereafter referred to as helix 26) and across the 3’-end of helix 45 through the D-site which generates the 3’-end of 18S after cleavage in the cytoplasm. Consistent Utp14-specific reads were also obtained at helices 18 and 36/37, while read densities across 21es6d (hereafter referred to as ES6) were reproducible but more variable between the two data sets. A small subset of reads aligned to nucleotides surrounding ∼480-600 of the 5’-ETS and the 5’-end of 18S. Mapping these binding sites to a current SSU Processome structure (Figure 1E) showed that helix 26 and the D-site are approximately 60 Å apart from one another, while helix 18 is tucked within the core of the structure, and the 5’-ETS sites are on the exterior of the particle approximately 70 Å away from helix 26 and approximately 140 Å away from the D-site. Helices 21es6d, 36, and 37 of the 18S rRNA were unresolved in this structure. These results imply that Utp14 traverses a large area of the SSU Processome. It is also possible that different sites are contacted at different stages of 40S assembly as Utp14 associates with 90S and pre-40S (Zhu et al. 2016).

Our result that Utp14 crosslinked across the A_1_ site and 5’-ETS is consistent with recent structures of the SSU Processome in which limited regions of Utp14 were resolved (Barandun et al. 2017). That work showed that residues 845-849 of Utp14 contact the A_1_ site and residues 828-834 of Utp14 contact several nucleotides of helix V of the 5’-ETS, while residues 317-408 and 876-896 of Utp14 wrap around helices VII and VIII of the 5’-ETS. A similar interaction of Utp14 with the 5’-ETS is also observed in the SSU Processome from the thermophilic fungus *Chaetomium thermophilum* (Cheng et al. 2017).

Since Utp14-HTP also enriched for snoRNAs (Figure 1B), we analyzed the percentage of reads aligning to each snoRNA relative to the total sense aligned reads (Supplemental File 1). U3 snoRNA (*SNR17A/B*) was present in both datasets with the majority of the reads mapping to nucleotides ∼20-60 of U3 (Figure 1F). Interestingly, this binding site overlaps the binding site of Dhr1 on U3 that we previously identified (Figure 1E) (Sardana et al. 2015). Although the negative control from data set 2 also contained reads to this region of U3, a recent SSU Processome structure confirmed that Utp14 appears to contact U24 and G37 of U3 (Barandun et al. 2017) which are within the range protected by Utp14 in our crosslinking analysis (Figure 1F). Thus, in addition to its rRNA contacts, we conclude that Utp14 also directly interacts with the U3 snoRNA. Moreover, a small set of reads aligned to snR30. It was previously reported that snR30 hybridizes to helix 26, a major crosslink site of Utp14, before its release by Rok1 (Martin et al. 2014). Thus, the reads mapping to snR30 may reflect a transient interaction between Utp14 and snR30. When taken together, these data demonstrate that Utp14 is an RNA binding protein that contacts multiple RNA elements within the SSU Processome with its primary sites being helix 26 and the D-site.

### The N-terminus of Utp14 interacts with proteins that bind Helix 26

We first attempted to support our crosslinking result that Utp14 binds to helix 26 using the yeast three-hybrid system, but we were unable to detect a specific interaction (data not shown). As an alternative approach, we reasoned that Utp14 may interact with proteins in the vicinity of its RNA binding sites. Utp22, Rrp7, and Rps1 (eS1) are within close proximity of helix 26 in recently solved structures of the SSU Processome (Sun et al. 2017; Cheng et al. 2017) (Figure 2A). Utp22 and Rrp7 are components of the UTPC sub-complex that are recruited to the pre-ribosome after synthesis of the Central domain of 18S (Chaker-Margot et al. 2015; Zhang et al. 2016; Lin et al. 2013). Additionally, an interaction between Utp22 and Utp14 was recently reported in two large-scale yeast two hybrid (Y2H) analyses of ribosome biogenesis factors (Baßler et al. 2016; Vincent et al. 2018). Rps1 is an r-protein needed upstream of processing at A_0_, A_1_, A_2_, and D (Ferreira-Cerca et al. 2005), and remains bound to helix 26 in the mature 40S (Ben-Shem et al. 2011). The TPR domain repeats of Rrp5 also bind near helix 26 the TPR domain repeats of Rrp5 (Sun et al. 2017), and an interaction between *C. thermophilum (ct)* Utp14 and Rrp5 was recently reported (Baßler et al. 2016).

**Figure 2.**
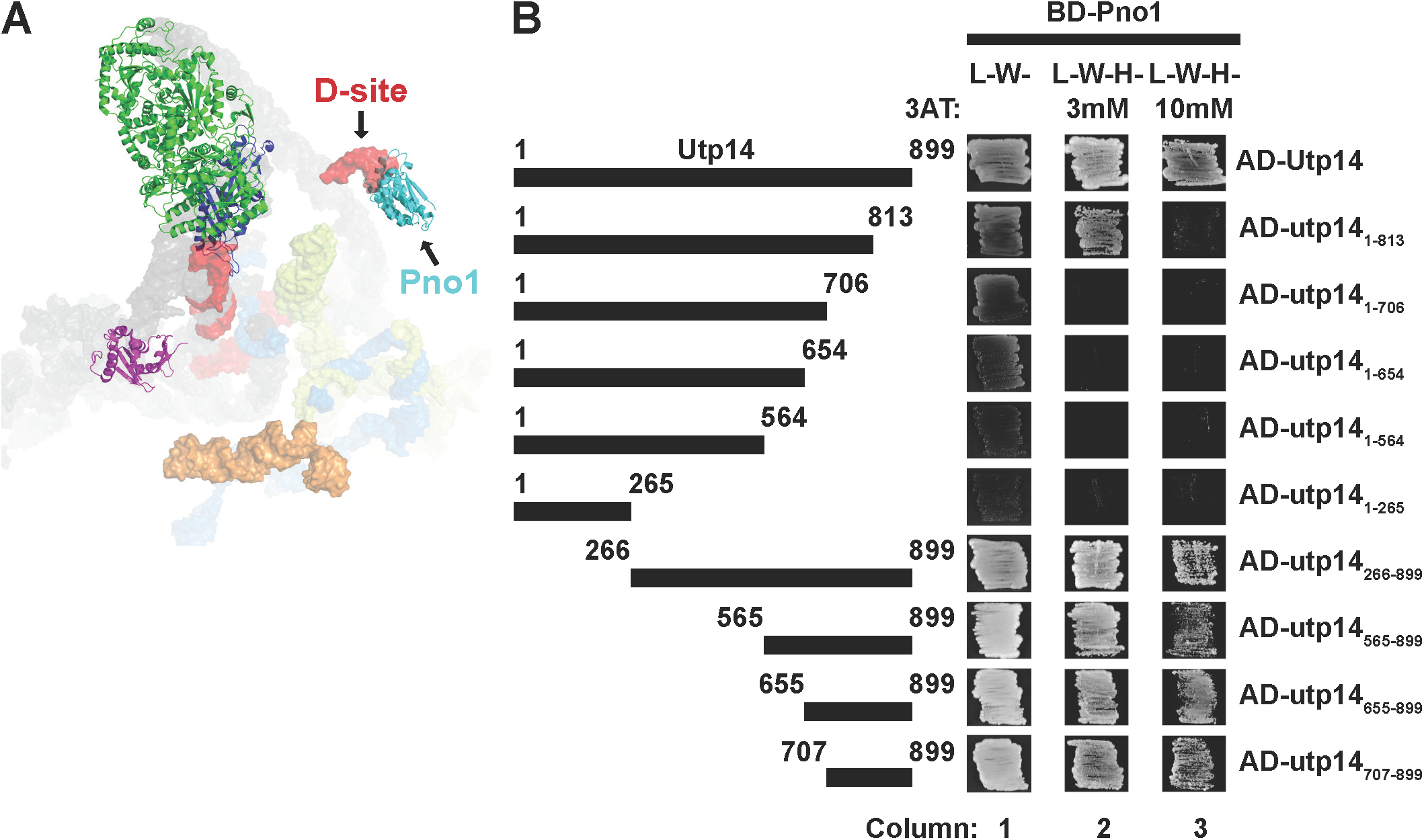
Residues 1-265 of Utp14 interact with proteins associated with helix 26. (A) Proteins that bind near Utp14 crosslinking sites in the SSU Processome (PDB: 5WYJ). Utp22 (green), Rps1 (blue), and Rps7 (magenta) are shown. Pno1 (cyan) is also shown for perspective. Utp14 crosslinking sites are shown in red (18S rRNA) and orange (5’-ETS). 18S rRNA (grey), 5’-ETS (yellow), and U3 (light blue) is shown in surface representation. (B) Yeast two-hybrid interaction data between Utp14 and Utp22 and Utp14 and Rps1 are shown. Strains carrying the indicated constructs were patched onto Leu^−^ Trp^−^ (L^−^W^−^) and Leu^−^ Trp^−^ His^−^ (L^−^W^−^H^−^) media supplemented with 3-Amino-1,2,4-triazole (3AT) as indicated. (BD, GAL4BD; AD, GAL4AD). A cartoon of the Utp14 constructs indicating amino acid positions is shown to the right.

We used Y2H analysis to test direct interactions between Utp14 and these proteins. Indeed, full length Utp14 interacted with Utp22 and Rps1 as indicated by growth on reporter media containing 3-amino-1,2,4-triazole (3AT), a competitive inhibitor of the *HIS3* gene product that increases the stringency of the assay. (Figure 2B; see columns 2 and 4). Utp14 is 899 amino acids in length, and much of the protein is not resolved in recent SSU Processome structures. To determine which region of Utp14 interacts with these proteins, we assayed a series of N- and C-terminal truncations of Utp14 for interaction. All C-terminal truncations retained interaction with Utp22 and Rps1. In contrast, all N-terminal deletions of Utp14 that were tested lost interaction with Utp22 and Rps1. Thus, the N-terminal portion (residues 1-265) of Utp14 was both necessary and sufficient for interaction with Utp22 and Rps1. We also noted that the interaction between Utp14 and Rps1 was enhanced by deletion of aa 565 to 899 (Figure 2B; *cf.* columns 4 and 5). The longer fragments of the protein may fold in a way that inhibits their interaction with Rps1 outside the context of the SSU Processome. We did not detect interactions between Utp14 and Rrp5 or Rrp7 using the *S. cerevisiae* genes (data not shown). Taken together, the Y2H interaction data between Utp22 and Rps1 with Up14 and the UV crosslinking of Utp14 to helix 26 suggests that the N-terminus of Utp14 (residues 1-265) is responsible for its interaction with helix 26.

### A C-terminal portion of Utp14 interacts with Pno1

To support the Utp14 crosslinks mapping to the D-site, we also first tested the interaction between Utp14 and the D-site by yeast three-hybrid but were unable to detect an interaction (data not shown). Consequently, we again considered that Utp14 may interact with proteins in the vicinity of the D-site. Recent structures of the SSU Processome show that Pno1 (Dim2) binds the D-site (Barandun et al. 2017; Sun et al. 2017) (Figure 3A). Pno1 is an essential KH-like domain protein that stably associates with the pre-ribosome once the majority of the 3’-minor rRNA domain of 18S is synthesized (Zhang et al. 2016; Chaker-Margot et al. 2015) and remains on pre-40S particles that enter the cytoplasm (Vanrobays et al. 2004; Johnson et al. 2017; Heuer et al. 2017). Pno1 is thought to recruit the dimethyltransferase Dim1, that methylates the 3’-end of the 18S rRNA (Vanrobays et al. 2004). An interaction between *ct*Utp14 and *ct*Pno1 was also reported in a large-scale screen for interactions among biogenesis factors (Baßler et al. 2016). To define the domain of Utp14 that interacts with Pno1, we again used Y2H analysis to assay interactions between the various Utp14 fragments and Pno1 (Figure 3B). Due to the proximity of the D site to helix 26, we initially expected Pno1 to interact with the N-terminus of Utp14. However, we found that C-terminal truncations abolished or weakened the interaction of Utp14 with Pno1, while N-terminal truncations maintained the interaction (Figure 2C; column 3). Furthermore, the Utp14 fragment containing residues 1-813 maintains an interaction with Pno1 on 3mM 3AT (Figure 3B; column 2) suggesting that residues 707-813 of Utp14 are critical for the interaction with Pno1. While this manuscript was in preparation an interaction between *ct*Utp14 and the KH-like domain of *ct*Pno1 was reported (Sturm et al. 2017). Taken together with that study, we infer that Utp14 binds to or near the D-site, and that the binding interface required for this interaction is between the KH-like domain of Pno1 and residues 707-813 of Utp14.

**Figure 3.**
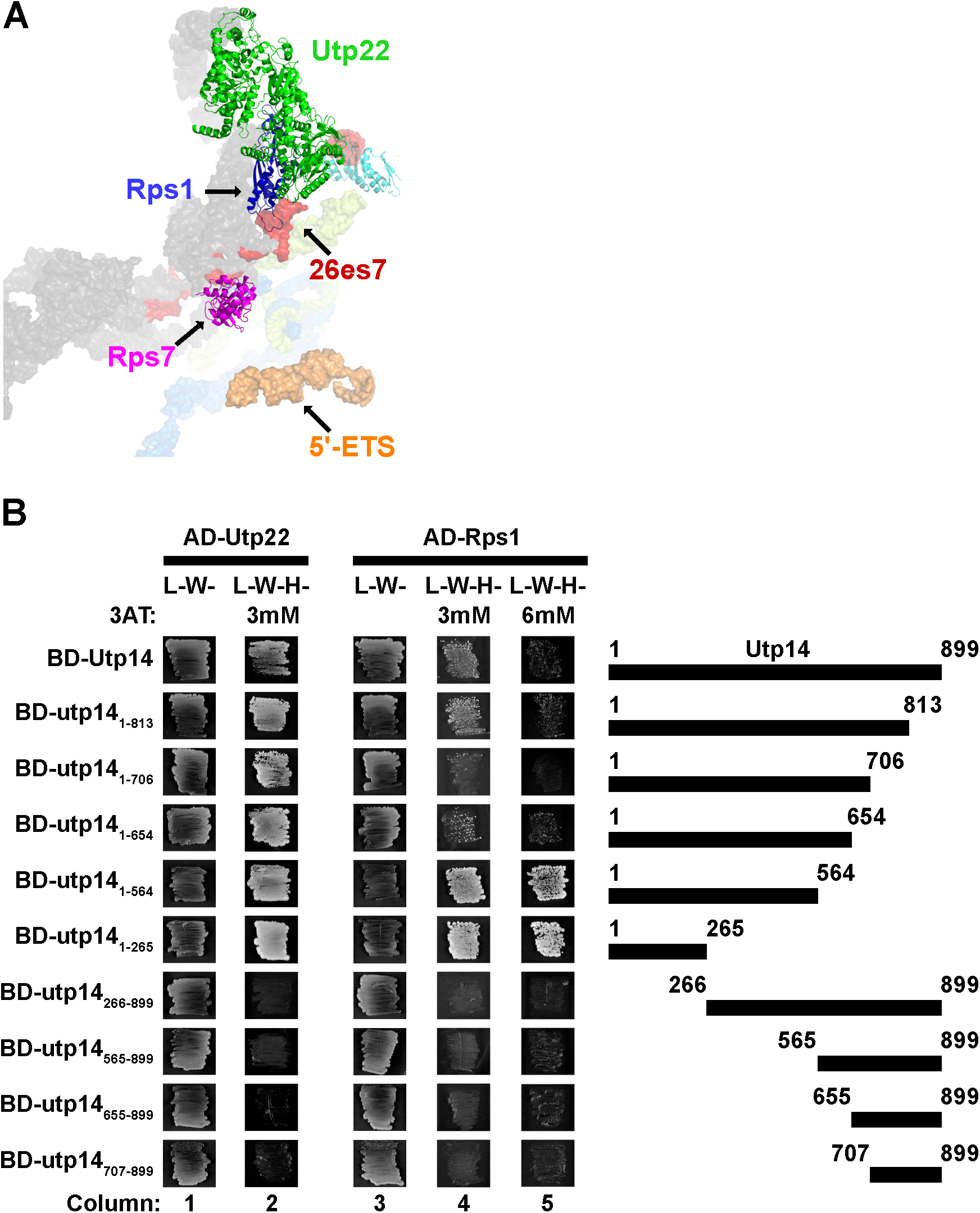
Residues 707-813 of Utp14 interact with Pno1. (A) Proteins that bind in the vicinity of the Utp14 crosslinking sites in the SSU Processome (PDB: 5WYJ). Coloring is the same as Figure 2A. (B) Yeast two-hybrid interaction data between Utp14 fragments and Pno1 are shown. Strains carrying the indicated constructs were patched onto Leu- Trp- (L-W-) and Leu- Trp- His- (L-W-H-) media supplemented with 3AT. (Abbreviations as used in Fig. 2) A cartoon of the Utp14 constructs indicating amino acid positions is shown to the left.

### Protein composition of wild-type and mutant Utp14 particles

Since Utp14 interacts with multiple regions of the SSU Processome, we subsequently sought to further understand how the presence of Utp14 affects the proteomic composition of preribosomal particles. We previously showed that Utp14 interacts with and activates the RNA helicase Dhr1 (Zhu et al. 2016). Both proteins are recruited to the pre-ribosome at a similar stage of maturation, (Chaker-Margot et al. 2015; Zhang et al. 2016) and thus are expected to stall progression of the SSU Processome at a similar point. To ask if Utp14 is required for the recruitment of additional proteins, we compared the protein composition of particles depleted of Utp14 or Dhr1. We isolated pre-ribosomal particles from cultures expressing C-terminally tagged tandem affinity purification (TAP) Enp1 after the repression of transcription of *UTP14* or *DHR1* and depletion of the respective proteins. Enp1 is an ideal bait for this assay as it binds prior to Utp14 association with pre-ribosomes (Zhang et al. 2016) and remains associated with pre-40S particles until the cytoplasm (Johnson et al. 2017), after Utp14 has been released. After affinity-purification and TEV elution, we sedimented samples through sucrose cushions to separate pre-ribosomal particles from extraribosomal bait and other co-purifying extraribosomal proteins. Following mass spectrometry, we generated relative spectral abundance factor (RSAF) values as described previously (Sardana et al. 2015). Figure 4A shows a heat map of RSAF values for 40S biogenesis factors that co-purified with the Enp1-TAP particles, normalized to the mean RSAF value for the UTP-B sub-complex of the sample as done previously (Zhang et al. 2016). This semi-quantitative analysis reflected the relative stoichiometry of proteins within the purified particles, validated by the 2-fold abundance of factors known to be present as dimers (Emg1 and Kre33) or in a 2:1 stoichiometry (Nop1 and Snu13) (Sun et al. 2017; Barandun et al. 2017; Cheng et al. 2017).

**Figure 4.**
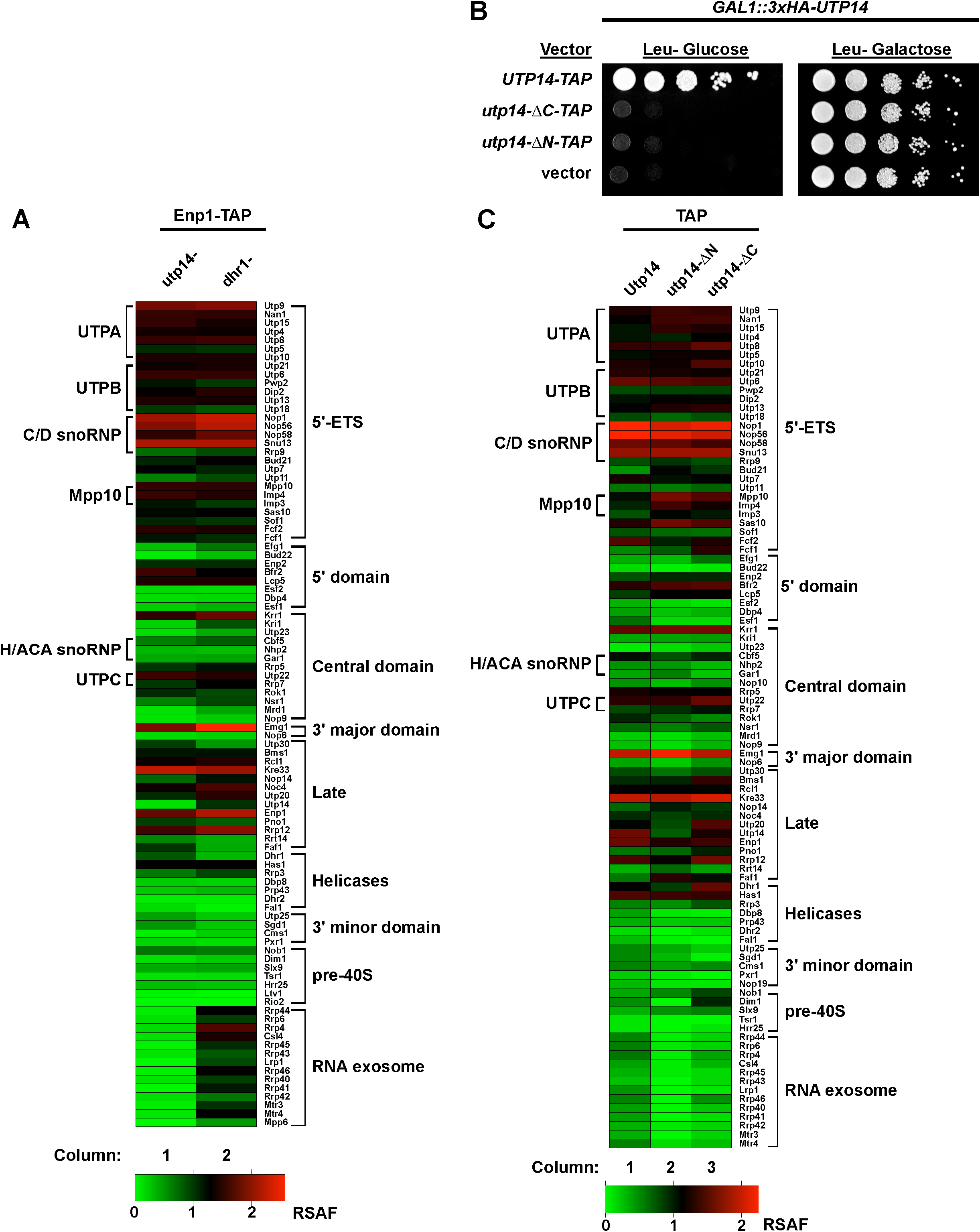
Proteomic profiles of mutant Utp14 particles. A heatmap representing the RSAF of each assembly factor identified by mass spectrometry relative to the mean RSAF of the UTP-B sub-complex of each sample for the affinity-purifications of (A) the Enp1-TAP constructs from strains depleted of Utp14 or Dhr1. (B) Ten-fold serial dilutions of AJY3243 cells harboring the indicated plasmids were grown at 30°C for two days. (C) A heatmap displaying data processed in the same manner as (A) but from strains harboring the Utp14-TAP constructs. Coloring for the heatmaps reflect apparent stoichiometry of each factor: green color represents RSAF values less than zero, black color represents an RSAF value of approximately one, and red color represents an RSAF of approximately two or greater. Proteins are grouped according to their order of recruitment to the pre-ribosome as reported in (Zhang et al. 2016) or by function. Heatmaps were generated in Graphpad Prism version 7.0c.169 for Mac iOS (www.graphpad.com). The data these heatmaps represent are reported in Supplemental File 2.

The overall compositions of the Utp14- and Dhr1-depleted particles were similar (Figure 4A and Supplemental File 2), however, we note several differences amongst the particles after applying the criteria that a given factor in the truncated Utp14 particles showed a log_2_-fold change of ± 1 or more relative to the Dhr1-depleted particles and at least one of the samples contained at least 5 spectral counts. As expected the Utp14-and Dhr1-depleted particles showed significantly reduced signal for Utp14 and Dhr1, respectively. The most notable difference between these two particles was a strong reduction of the RNA exosome in the Utp14-depleted particle relative to the Dhr1-depleted particle suggesting that Utp14 may have a role in the recruitment of the exosome.

As a complementary approach, we also tested the significance of the interactions of the N- and C-terminal regions of Utp14 using two truncation mutants deleted of residues 1-265 (Utp14- ΔN) or residues 707-899 (Utp14-ΔC). These Utp14 mutants are expected to lose interactions with its binding sites at helix 26 and the D-site, respectively (Figures 2 and 3). Neither of these truncation mutants was able to complement the loss of Utp14 (Figure 4B). To rule out the possibility that these mutants were non-functional because they failed to express well or engage with pre-ribosomes, we assayed their sedimentation in sucrose gradients. Both the N- and C-terminally truncated proteins showed a population of protein that sedimented in the 40S to 80S region of the gradient, similar to wild-type, suggesting that both proteins enter into pre-ribosomal particles (data not shown). However, less Utp14-ΔC sedimented in these deeper gradient fractions than Utp14-ΔN did, suggesting that the association of Utp14-ΔC with pre-ribosomes is reduced.

To ask if there were any changes in the protein composition of the Utp14 mutant particles compared to wild-type, we affinity-purified particles via C-terminal TAP tags. The particles affinity-purified by wild-type or truncated Utp14 displayed overall similar protein compositions with some differences (Figure 4C and Supplemental File 2). Most notably, the full-length Utp14 particles contained the exosome, while it was nearly absent from the Utp14-ΔN, and significantly reduced in the Utp14-ΔC particle further suggesting that Utp14 has a role in the efficient recruitment of the exosome. Moreover, the Utp14-ΔC particles were enriched for Nob1 and slightly enriched for Dim1 and its interacting partner Pno1. Conversely, the Utp14-ΔN particles completely lacked Dim1 but contained wild-type levels both of Nob1 and Pno1. In general, the Utp14-ΔC particles contained a greater abundance of pre-40S factors than the full-length Utp14 or Utp14-ΔN particles. These observations support the notion that the Utp14-ΔN particle is stalled in the SSU Processome assembly pathway upstream of the Utp14-ΔC particle. Furthermore, both mutant particles displayed overall reduced signal relative to the full length Utp14 particles for the RNA helicases including Dbp8 and its cofactor Esf2 and overall decrease in 3’ minor domain factors Utp25 and Sgd1 suggesting that the Utp14 mutants have an effect on the recruitment of these factors. Together, the results of both analyses suggest an unexpected role for Utp14 in the recruitment of the exosome to the SSU Processome.

### Utp14-ΔC co-purifies with an extraribosomal sub-complex containing Rps7 and Rps22

Our purification strategy for pre-ribosomal particles, involving sedimentation of particles through a sucrose cushion, separated bait associated with pre-ribosomes from extraribosomal bait and other non-ribosome-bound proteins. We noted two lower molecular weight species present in the Utp14-ΔC-TAP extraribosomal fraction (Figure 5A; lane 6). Mass spectrometry identified the ∼ 20 kDa species to be Rps7 (eS7) and the ∼10 kDa species to be Rps22 (uS8). Rps7 and Rps22 interact directly with one another in the context of nascent and mature ribosomes (Figure 5B) (Ben-Shem et al. 2011; Sun et al. 2017; Barandun et al. 2017; Cheng et al. 2017) and we recapitulated this interaction by Y2H (Figure 5C). We then used the Y2H system to ask if Utp14 interacted with either Rps7 or Rps22. We found that the N-terminus of Utp14 (residues 1-265) was both necessary and sufficient for the interaction between Utp14 and Rps7 (Figure 5D). We did not, however, detect an interaction between Utp14 and Rps22 (data not shown). These results suggest that Rps7 and Rps22 initially bind to the SSU Processome in an unstable fashion and require full length Utp14 to stabilize their interaction with the SSU Processome. We considered the possibility that Rps7 or Rps22 is needed for the recruitment of Utp14 to the SSU Processome but did not observe any decreased association of Utp14 upon depletion of either Rps7 or Rps77 (data not shown).

**Figure 5.**
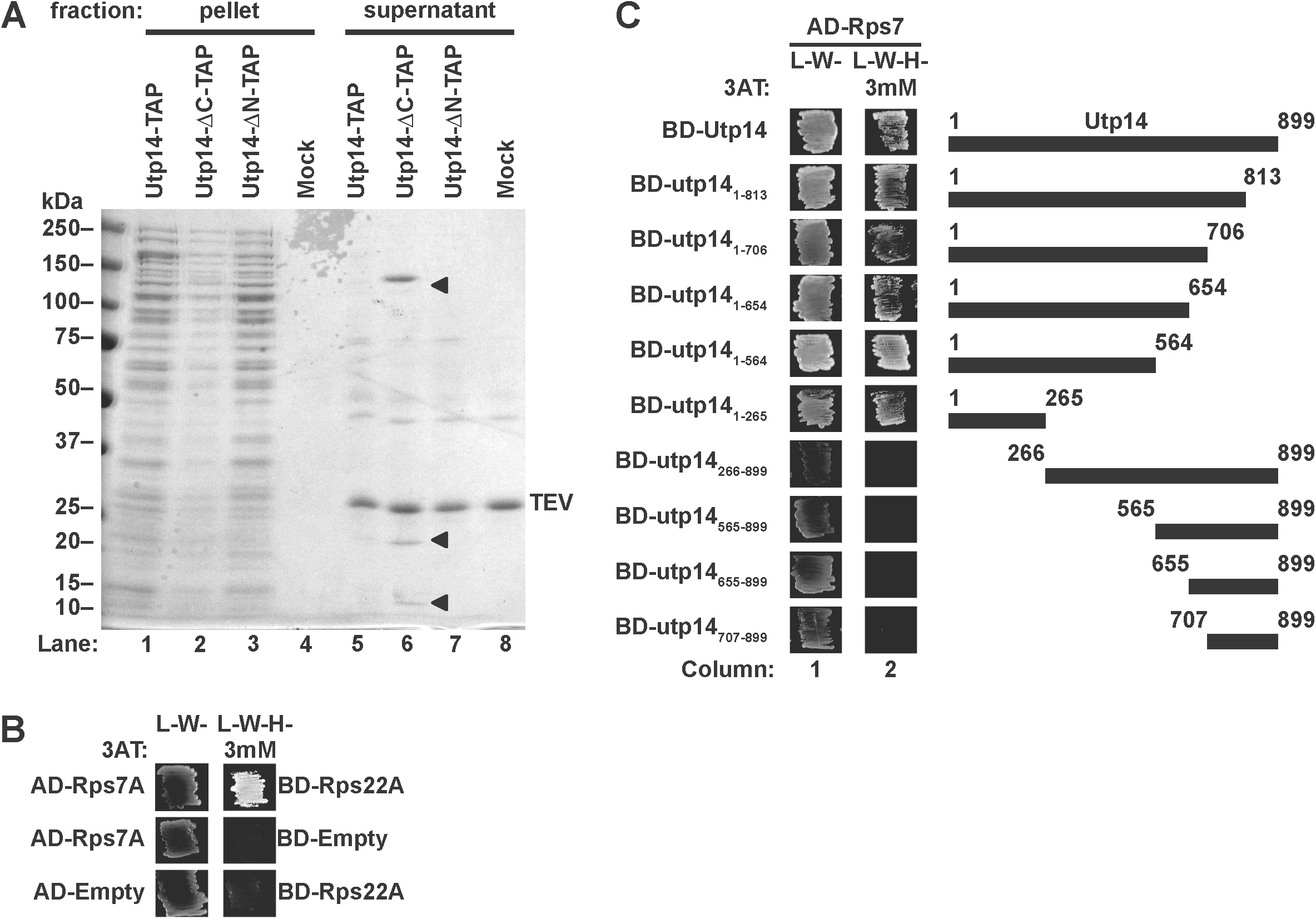
The N-terminus of Utp14 interacts with an extraribosomal Rps7-Rps22 heterodimer. (A) Coomassie-stained gel of proteins that co-purified with full-length or truncated Utp14 proteins. Pellet and supernatant fractions were separated by overlaying eluate onto sucrose cushions followed by ultracentrifugation. The arrow heads in lane 6 indicate Utp14-ΔC (∼150kDa), Rps7 (∼20kDa), and Rps22 (∼10kDa). (B) Yeast two-hybrid interaction data for Rps7 and Rps22. (C) Yeast two-hybrid interaction data between Utp14 and Rps7 are shown. A cartoon of the Utp14 constructs is shown to the right. Abbreviations as in legend of Fig. 2.

### RNA composition of wild-type and mutant Utp14 particles

We next asked whether the Utp14 mutant particles also differed in their content of rRNA processing intermediates. RNA was prepared from TAP-tagged wild-type and Utp14 mutant particles and analyzed by northern blotting to detect rRNA processing intermediates (Figure 6A). Wild-type and truncated mutant Utp14 associated with distinct rRNA processing intermediates consistent with their ability to bind pre-ribosomal particles (Figure 6B). Particles pulled down with full-length Utp14 contained 35S, 33S, 23S, 22/21S, and 20S rRNA intermediates (Figure 6B; lane 1), reflecting its association with the SSU processome and pre-40S at multiple stages of pre-rRNA processing. Utp14-ΔN associated with 35S, 33S, 23S, and 22/21S, but not 20S (Figure 6B; lane 2, D-A_2_ panel). The Utp14-ΔN particle was also enriched for degradation intermediates of 23S rRNA (asterisks in Fig 6B), suggesting that the associated RNA was subjected to 3’-degradation by the exosome. Similar to full length Utp14, the Utp14-ΔC mutant associated with 35S, 33S, 23S, 22/21S, and 20S (Figure 6B; lane 3), but co-purified with less rRNA overall, consistent with its decreased association with the pre-ribosome (Figure 5A). The lack of 20S rRNA in the Utp14-ΔN particle, indicates that this particle is stalled earlier in the processing pathway, at A_2_ cleavage, compared to the Utp14-ΔC mutant particle. This result agrees with the proteomic profiles described above, in which the Utp14-ΔN contained overall less pre-40S factors than the Utp14-ΔC (Figure 4A).

**Figure 6.**
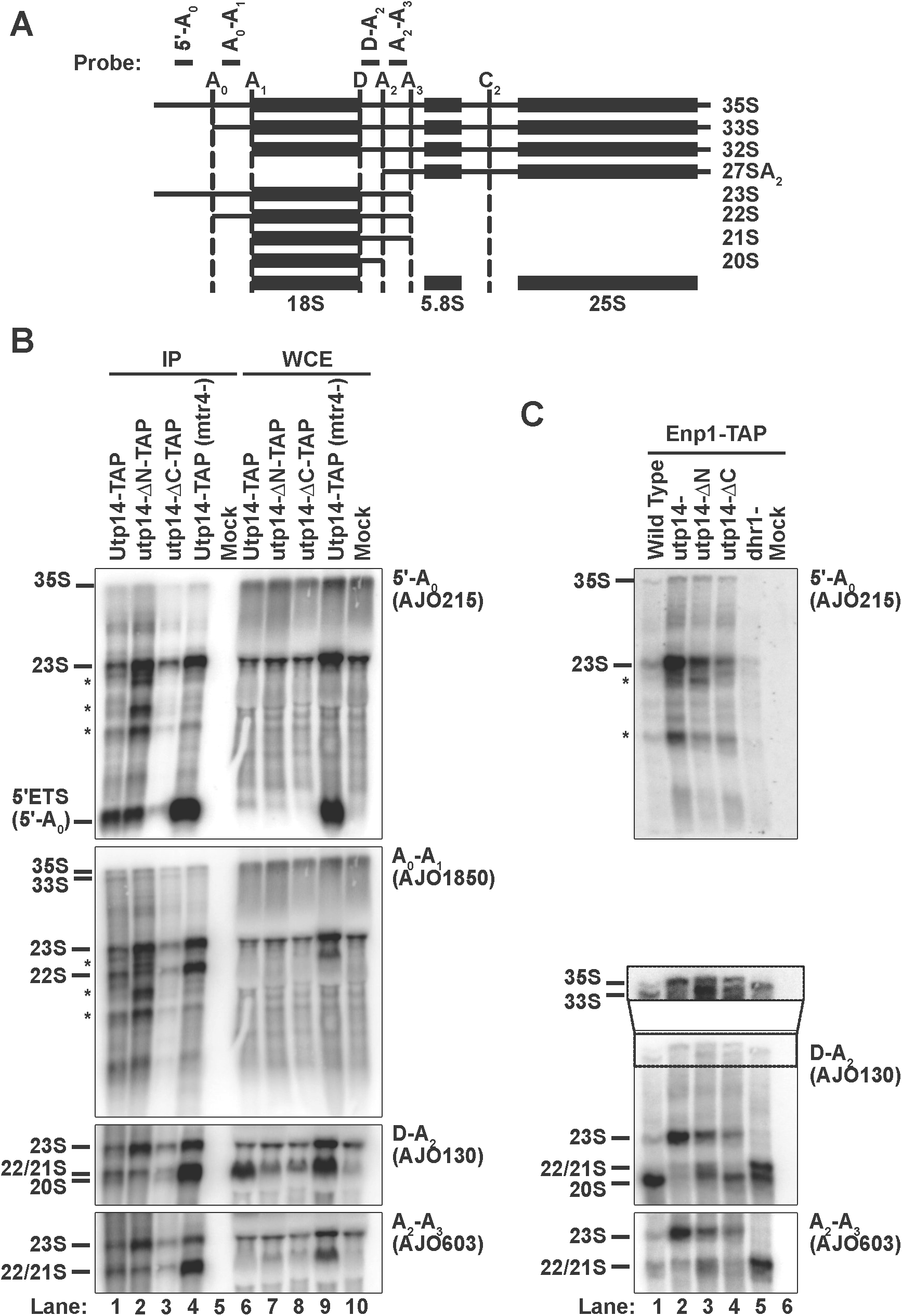
The rRNA processing intermediates associated with mutant Utp14 particles. (A) A cartoon of rRNA processing and oligos used to detect intermediates. The sequences for the probes are:5’-A_0_(5’-GGTCTCTCTGCTGCCGGAAATG-3’),A_0_-A_1_ (5’-CCCACCTATTCCCTCTTGC-3’), D-A_2_ (5’-TCTTGCCCAGTAAAAGCTCTCATGC-3’), A_2_-A_3_ (5’-TGTTACCTCTGGGCCCCGATTG-3’). (B, C) Northern blots for rRNA processing intermediates affinity-purified via (B) TAP-tagged Utp14 full length and truncation mutants or untagged wild-type (mock) and (C) TAP-tagged Enp1 from cells depleted of Utp14 or Dhr1, conditionally expressing Utp14 truncation mutants or from untagged Enp1 (mock). Images were captured on a Typhoon FLA9500 and processed in ImageJ.

The exosome is required for the exonucleolytic degradation of the 5’-A_0_ fragment (Thoms et al. 2015). Because the mutant Utp14 particles were deficient for the exosome (Figure 4A), we asked if the 5’-A_0_ fragment was enriched in the Utp14 mutant particles. For comparison, we depleted the exosome-associated helicase Mtr4 (Figure 6B; lane 4) to inhibit degradation of the 5’-A_0_ fragment (Thoms et al. 2015). The Mtr4-depleted sample was highly enriched for the 5’-A_0_ fragment, as expected. However, the Utp14-depleted particle did not show a similar enrichment for this fragment despite being severely depleted for the exosome, suggesting that the apparent lack of exosome recruitment in the Utp14 mutant particles does not result in a noticeable defect in degradation of 5’-A_0_ in these particles.

As an alternative method to ask how the Utp14 mutants affected rRNA processing, we carried out a second set of purifications using Enp1-TAP from cells expressing wild-type Utp14, the N- and C-terminal truncation mutants or depleted of Utp14. For comparison, we also affinity purified Enp1-TAP from Dhr1-depleted cells. Northern blot analysis of the RNAs that co-purified with Enp1-TAP from wild-type cells revealed that Enp1 primarily associated with 20S but low levels of 33S, 23S and 22S/21S were also observed (Figure 6C; lane 1). This result is consistent with the late entry of Enp1 into the SSU Processome and its continued association with pre-40S (Chaker-Margot et al. 2015; Zhang et al. 2016; Johnson et al. 2017; Sun et al. 2017). In the absence of Utp14, Enp1-TAP associated primarily with 23S (Figure 6C; lane 2) with a low level of 22S/21S also detected. Interestingly, in the absence of Utp14, Enp1 also associated with low levels of 35S but not 33S as observed in wild-type cells. Apparently, in the absence of Utp14, cleavage at A_0_ is blocked and Enp1 is recruited to 35S instead of 33S. The strong accumulation of 23S suggests that Utp14 is also required for cleavages at A_1_ and A_2_. The low level of 22S/21S may be due to continued processing in the presence of residual Utp14 or indicate that Utp14 is not absolutely required for A_1_ and A_2_ cleavage.

The two Utp14 truncation mutants resulted in Enp1 association with RNAs reflecting processing that was intermediate between that of wild-type and Utp14-depleted cells. In the presence of Utp14-ΔN, Enp1 associated with both 35S and 33S RNAs and instead of the strong accumulation of 23S in Utp14-depleted cells or 20S in wild-type cells, signal was roughly equally distributed among 23S, 22/21S, and 20S species (Figure 6C; lane 3). In contrast, in the presence of Utp14-ΔC Enp1 associated with both 35S and 33S but the levels of 23S and 22S/21S were reduced with 20S predominating (Figure 6C; lane 4). The presence of 20S in the Enp1-purified particle from Utp14-ΔN-expressing cells was surprising given that Utp14-ΔN itself does not co-purify with 20S (Fig. 6B; lane 2). This may indicate that while Utp14-ΔN associates with pre-ribosomes that have not yet been cleaved at A_2_, it may not stably associate with particles after A_2_ cleavage. These results indicate that the N- and C-terminally truncated proteins support rRNA processing that is intermediate between that of wild-type and Utp14-depleted cells, with the Utp14-ΔC mutant supporting more extensive processing. By comparison, in the absence of Dhr1 Enp1 co-purified with 33S, 22/21S and 20S but not 35S or 23S (Figure 6C; lane 5), indicating that Utp14 is required upstream of Dhr1 for cleavages at A_0_ and A_1_. The accumulation of 22/21S rRNA from Utp14-ΔN-expressing cells was similar to the processing defects of the Dhr1-depleted particles, suggesting the Utp14-ΔN is defective in its ability to stimulate Dhr1 efficiently (Figure 6C; *cf.* lanes 3 and 5).

## DISCUSSION

We previously identified Dhr1 as the RNA helicase that unwinds U3 from the pre-rRNA (Sardana et al. 2015). Considering the central role that U3 hybridization to the pre-rRNA plays in organizing the structure of the SSU Processome, its unwinding by Dhr1 likely contributes to disassembly of the SSU Processome in the transition to the pre-40S particle. What times the activation of Dhr1, to unwind U3 at the appropriate stage of SSU Processome assembly remains an open question. We identified Utp14 as a Dhr1-interacting partner that stimulates the unwinding activity of Dhr1 (Zhu et al. 2016), raising the possibility that Utp14 is involved in timing Dhr1 activity *in vivo*. In an effort to understand how Utp14 might coordinate SSU Processome assembly with stimulation of Dhr1 activity, we mapped the interaction of Utp14 with the pre-ribosome identifying that Utp14 binds to multiple regions within the pre-18S rRNA, including 5’ and 3’ elements. While this manuscript was in preparation, the partial structure of Utp14 in the SSU Processome was solved (Barandun et al. 2017; Cheng et al. 2017). Our analysis extends the structural analysis by uncovering how unresolved elements of Utp14 interact with the pre-ribosome. Moreover, our analysis suggests a model in which Utp14 communicates between the 5’- and 3’-ends of pre-18S rRNA to monitor the status of the SSU Processome (see below). Our proteomic characterization of pre-ribosomal particles depleted of Utp14 revealed a specific loss of the exosome, responsible for the exonucleolytic degradation of the 5’-ETS. These results could suggest an unanticipated role for Utp14 in the recruitment of this complex.

### Does Utp14 communicate between the 3’- and 5’-ends of 18S rRNA?

Our protein-RNA crosslinking analysis identified a major binding site for Utp14 across the D-site of pre-rRNA, the cleavage site that generates the mature 3’-end of 18S. We also identified Utp14 binding sites within the 5’-ETS and across the A_1_ site, which generates the mature 5’-end of 18S (Figure 1 C, D). To complement our UV crosslinking approach, which did not allow us to determine the domains of Utp14 that were responsible for these RNA interactions, we used yeast-two hybrid analysis to identify interactions between domains of Utp14 and proteins that bound in the vicinity of the RNA binding sites, thereby approximating the domains of Utp14 responsible for the major RNA interactions at helix 26 and the D site. In addition, several *α* - helices of Utp14 were assigned in recent cryo-EM structures of the SSU Processome (Barandun et al. 2017; Cheng et al. 2017), corroborating the interactions of Utp14 that we identified with the 5’-ETS and Site A_1_ and identifying the residues of Utp14 that are likely involved in these interactions. Because Utp14 associates with 35S pre-rRNA in the SSU Processome as well as 20S pre-rRNA in the pre-40S (Zhu et al. 2016), the RNA interactions we identified could reflect interactions at various stages of 40S biogenesis. However, the ability to map interactions to the SSU Processome suggests that the interactions we detected are predominantly in the context of the intact SSU Processome. These results are summarized in Figure 7, which combines our RNA crosslinking and protein interaction data with SSU Processome structure. We see that a C-terminal region of Utp14, between aa 707 and 813, interacts with the D site and Pno1 whereas the A_1_ site and 5’-ETS is recognized by a complex interaction of the extreme C-terminus of Utp14 and overlapping helices that wrap around the 5’-ETS. Connecting these two regions is a long unresolved loop that contains the Dhr1 binding site, from aa 565-813. Thus, Utp14 is uniquely positioned to connect the 5’- and 3’-ends of the 18S rRNA, tethering Dhr1 via the intervening loop. A tempting model is that Utp14 actively monitors the status of transcription and assembly of the 3’-end of the small subunit RNA, to signal maturation of SSU Processome. Such a model affords a mechanism for how Utp14 could time the of activation the helicase activity of Dhr1 to unwind U3. However, both Utp14 and Dhr1 are present in the mature SSU Processome, with U3 remaining bound to rRNA, indicating that additional signals are required to trigger Dhr1 unwinding.

**Figure 7.**
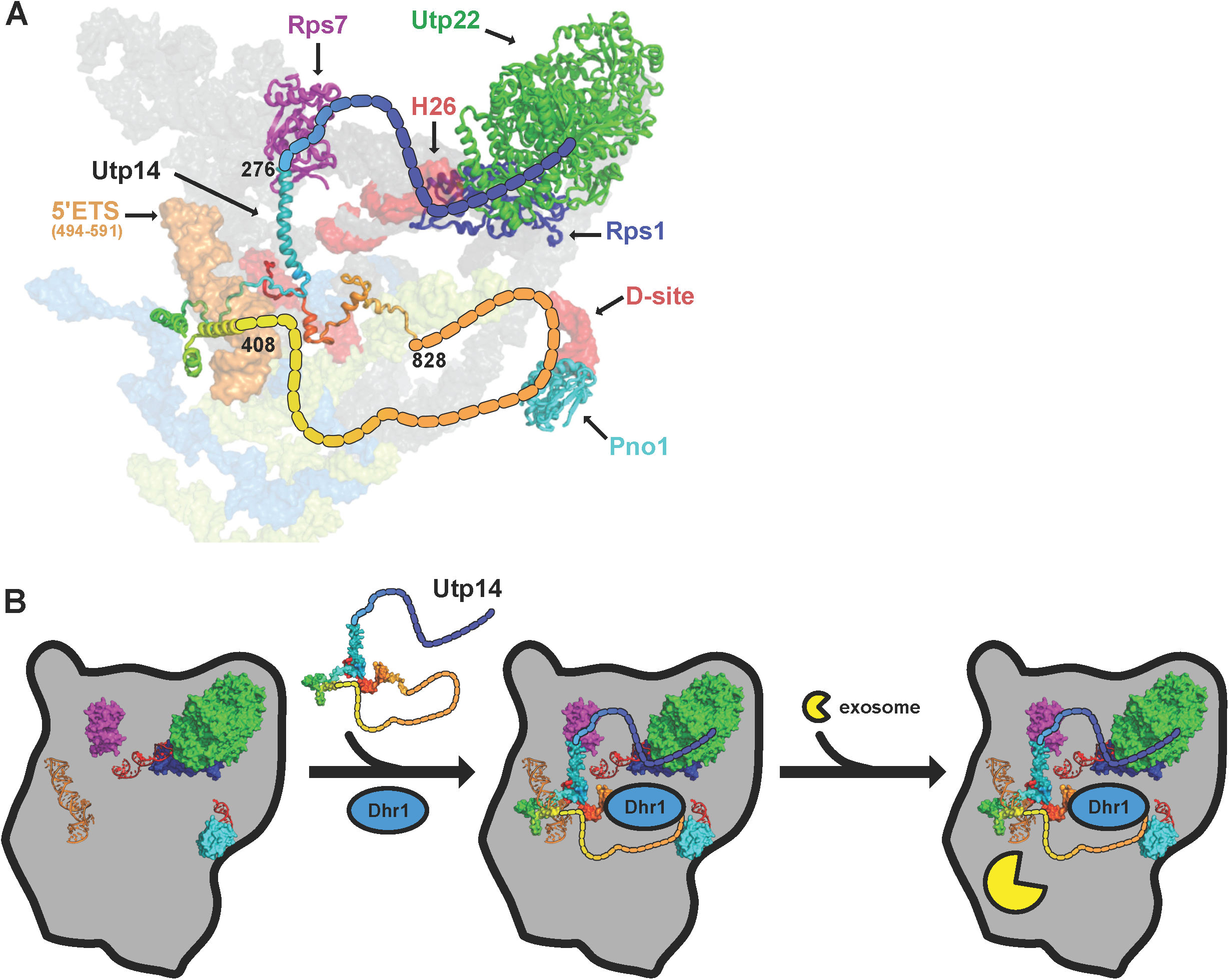
Model for Utp14 interaction with the SSU Processome. (A) A composite of Utp14 (rainbow; from PDB 5TZS) fitted into an SSU Processome structure (PDB 5WYJ) is shown. Dashed lines are shown to highlight the interactions identified in this work. Black numbers indicate the residues of Utp14 where the strands of resolved residues end. Rps7 (magenta), Rps22 (orange), Utp22 (green), Rps1 (blue), and Pno1 (cyan) are shown as cartoon representation. Binding site of Utp14 in 18S binding sites (red) and in 5’-ETS sites (orange), other regions of 18S rRNA (grey), 5-ETS (yellow), and U3 (light blue) are also shown as surface representation. (B) A model for the stepwise entry of Utp14, Dhr1, and exosome into the SSU Processome in which the assembly of Utp14 promotes the recruitment of the exosome. Relevant factors and rRNA elements are shown and colored the same as in (A).

### What is the relationship between Utp14 and the nuclear RNA exosome?

Our proteomic analysis of Utp14 mutant particles suggests that Utp14 is needed for the efficient recruitment of the exosome to the SSU Processome. This conclusion is based on our observation that the exosome was severely reduced in Utp14-depleted and mutant Utp14 particles compared with Utp14 replete or Dhr1-depleted particles. This difference in exosome abundance was despite the overall similarity in protein composition among these particles and suggests that either Utp14 is directly involved in recruiting the exosome or Utp14 is required for structural rearrangements of the SSU Processome that promote exosome recruitment.

It was previously shown that Utp18 recruits the exosome to the 5’-ETS through direct interaction between its AIM domain and the Arch domain of Mtr4, the RNA helicase for the nuclear exosome (Thoms et al. 2015). Utp18 is a component of the UTP-B sub complex and binds relatively early to the assembling nascent SSU Processome, after the 5’-ETS has been transcribed (Chaker-Margot et al. 2015; Zhang et al. 2016), but it is expected that the exosome is not recruited until the SSU Processome is fully assembled. To rationalize Utp18 recruitment significantly preceding exosome recruitment, it was proposed that accessibility of the AIM domain to Mtr4 is regulated during assembly of the SSU Processome (Thoms et al. 2015). It is possible that the recruitment of Utp14 regulates accessibility of the AIM domain. However, based on current structures in which limited portions of both Utp14 and Utp18 have been resolved, we prefer the model in which Utp14 is required together with Utp18 for stable recruitment of Mtr4. In those structures, residues near the N-terminus and the very C-terminus of Utp14 interact with each other, in the vicinity of the A_1_ site, the expected position of the A_0_ site and approaching Utp18. We note that the exosome was depleted from particles lacking Utp14 in its entirety or lacking either terminus. We suggest that interaction of the two termini is critical for establishing a structure that stabilizes the exosome, either through direct interaction or indirectly through the structure of the assembled SSU Processome. Such influence on the recruitment of the exosome would be consistent with the idea that Utp14 acts to signal between the 3’- and 5’-ends of the SSU Processome to recruit the exosome only after the SSU Processome has been deemed complete (Figure 7B).

## MATERIALS AND METHODS

### Strains, plasmids, and growth media

All *S. cerevisiae* strains and sources are listed in Table 1. AJY4051 was generated by genomic integration of the HIS6-tobacco etch virus (TEV)-protein A (HTP) tag (Granneman et al. 2009) into BY4741. AJY4257 and AJY4258 were generated by genomic integration of *ENP1-TAP::HIS3MX6* amplified from AJY2665 into AJY3243 and AJY3711, respectively. All yeast were cultured at 30°C in either YPD (2% peptone, 1% yeast extract, 2% dextrose), YPgal(2% peptone, 1% yeast extract, 1% galactose), or synthetic dropout (SD) medium containing 2% dextrose unless otherwise noted. All plasmids used in this study are listed in Table 2.

**Table 1:**
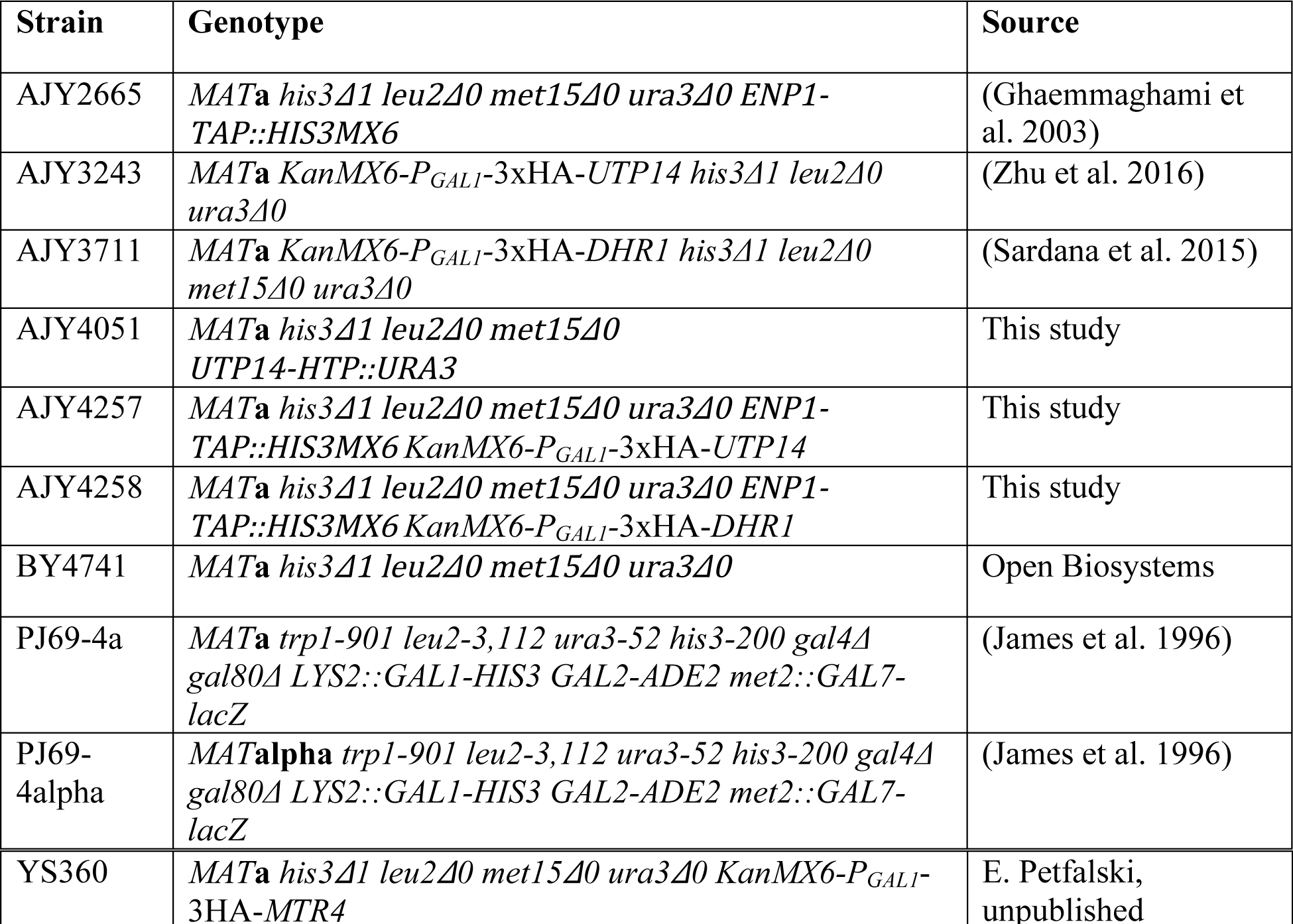
Strains used in this study.

**Table 2:** Plasmids used in this study.

| Plasmid | Description | Source |
| --- | --- | --- |
| pACT2 | GAL4AD-HA *LEU2* 2μ | Clontech |
| pAJ2321 | GAL4AD-HA-*UTP14* *LEU2* 2μ | (Zhu et al. 2016) |
| pAJ2324 | GAL4BD-c-myc-*UTP14* *TRP1* 2μ | This study |
| pAJ2334 | GAL4AD-HA-*utp14*<sub>1-706</sub> *LEU2* 2μ | (Zhu et al. 2016) |
| pAJ2335 | GAL4AD-HA-*utp14*<sub>707-899</sub> *LEU2* 2μ | (Zhu et al. 2016) |
| pAJ2341 | GAL4AD-HA-*utp14*<sub>1-813</sub> *LEU2* 2μ | (Zhu et al. 2016) |
| pAJ2342 | GAL4AD-HA-*utp14*<sub>1-654</sub> *LEU2* 2μ | (Zhu et al. 2016) |
| pAJ2343 | GAL4AD-HA-*utp14*<sub>1-564</sub> *LEU2* 2μ | (Zhu et al. 2016) |
| pAJ2344 | GAL4AD-HA-*utp14*<sub>1-265</sub> *LEU2* 2μ | (Zhu et al. 2016) |
| pAJ2345 | GAL4AD-HA-*utp14*<sub>266-899</sub> *LEU2* 2μ | (Zhu et al. 2016) |
| pAJ2346 | GAL4AD-HA-*utp14*<sub>565-899</sub> *LEU2* 2μ | (Zhu et al. 2016) |
| pAJ2347 | GAL4AD-HA-*utp14*<sub>655-899</sub> *LEU2* 2μ | (Zhu et al. 2016) |
| pAJ3422 | *utp14*<sub>1-706</sub> *URA3* *CEN* *ARS* | This study |
| pAJ3426 | *utp14*<sub>266-899</sub> *URA3* *CEN* *ARS* | This study |
| pAJ3624 | GAL4AD-HA-*RPS1A* *LEU2* 2μ | This study |
| pAJ3625 | GAL4BD-c-myc-*utp14*<sub>1-706</sub> *TRP1* 2μ | This study |
| pAJ3626 | GAL4BD-c-myc-*utp14*<sub>1-813</sub> *TRP1* 2μ | This study |
| pAJ3627 | GAL4BD-c-myc-*utp14*<sub>1-654</sub> *TRP1* 2μ | This study |
| pAJ3628 | GAL4BD-c-myc-*utp14*<sub>1-564</sub> *TRP1* 2μ | This study |
| pAJ3629 | GAL4BD-c-myc-*utp14*<sub>1-265</sub> *TRP1* 2μ | This study |
| pAJ3832 | GAL4BD-c-myc-*utp14*<sub>266-899</sub> *TRP1* 2μ | This study |
| pAJ3833 | GAL4BD-c-myc-*utp14*<sub>565-899</sub> *TRP1* 2μ | This study |
| pAJ3834 | GAL4BD-c-myc-*utp14*<sub>655-899</sub> *TRP1* 2μ | This study |
| pAJ3835 | GAL4BD-c-myc-*utp14*<sub>707-899</sub> *TRP1* 2μ | This study |
| pAJ3846 | GAL4AD-HA-*UTP22* *LEU2* 2μ | This study |
| pAJ4068 | GAL4BD-c-myc-*PNO1* *TRP1* 2μ | This study |
| pAJ4176 | *UTP14-TAP* *LEU2* *CEN ARS* | This study |
| pAJ4177 | *utp14*<sub>1-706</sub>-*TAP* *LEU2* *CEN ARS* | This study |
| pAJ4178 | *utp14*<sub>266-899</sub>-*TAP* *LEU2* *CEN ARS* | This study |
| pAJ4179 | GAL4AD-HA-*RPS7A* *LEU2* 2μ | This study |
| pAJ4182 | GAL4BD-c-myc-*RPS22A* *TRP1* 2μ | This study |
| pGADT7 | GAL4AD-HA *LEU2* 2μ | (Patel et al. 2007) |
| pGBKT7 | GAL4BD-c-myc *TRP1* 2μ | (Patel et al. 2007) |
| pRS415 | *LEU2* *CEN ARS* | (Sikorski and Hieter 1989) |
| pRS416 | *URA3* *CEN ARS* | (Sikorski and Hieter 1989) |

### Yeast two-hybrid (Y2H) analysis

GAL4 activation domain (AD)-containing vectors were transformed into PJ69-4a, and GAL4 DNA binding domain (BD)-containing vectors were transformed into PJ69-4alpha. Cells harboring these vectors were mated on YPD plates and then replica plated onto SD medium lacking leucine and tryptophan (SD Leu^−^ Trp^−^ medium) to select for diploid cells harboring both plasmids. The diploid strains were patched on SD Leu^−^ Trp^−^ and SD Leu^−^ Trp^−^ His^−^ with or without 3-amino-1,2,4-triazole (3AT) to test for activation of the *UAS*_*GAL*_-*HIS3* reporter gene.

### UV-<u>cr</u>osslinking and <u>a</u>nalysis of <u>c</u>DNA (CRAC)

A modified version of the CRAC protocol (Granneman et al. 2009) was performed. Cells from exponential phase cultures AJY4051 and BY4741 were collected, resuspended in PBS on ice and irradiated at 254nm using a Stratalinker UV Crosslinker 1800 with 800-1600 mJ/cm^2^ and stored at −80°C. Cells were resuspended in ice-cold TN150 buffer (50 mM Tris-HCl pH 7.8, 150 mM NaCl, 0.1% NP-40, 10 mM β-mercaptoethanol [BME], 2 mM MgCl_2_, 1 mM PMSF, and 1 µM leupeptin and pepstatin) and extracts were prepared by vortexing with glass beads, and clarified by centrifugation. Extract was incubated with IgG-Sepharose beads (GE Health Care) for 4 hours at 4°C. The beads were washed with ice-cold TN1000 buffer (TN150, except 1 M NaCl) then with ice-cold TN150 buffer lacking protease inhibitors. Protein was released from the resin using GST-TEV for 4 hours at 16°C with rotation. RNAs were digested with RNace-IT Ribonuclease Cocktail (Agilent Technologies) at 37°C. This mixture was then supplemented to final concentrations of 6 M guanidinium chloride, 10 mM imidazole, and 200 mM NaCl and bound to Ni-NTA resin (Invitrogen) overnight at 4°C.

The resin was washed with Buffer I (50 mM Tris-HCl pH 7.8, 300 mM NaCl, 10 mM imidazole, 6 M guanidinium chloride, 0.1% NP-40, and 10 mM BME) and with T4 Polynucleotide kinase (PNK) buffer (70 mM Tris-HCl pH 7.6, 10 mM MgCl_2_, and 10 mM BME). RNA retained on beads was labeled using T4 PNK and ^32^P-γ-ATP (PerkinElmer). T4 PNK also removes the 3’-phosphate remaining from RNase treatment. After labeling, the resin was washed with T4 PNK buffer, and AIR Adenylated Linker A (Bioo Scientific; 5’-rAppCTGTAGGCACCATCAAT/3ddC/-3’) was ligated at room temperature for 4-6 hours using T4 RNA Ligase 2 (truncated) (New England Biolabs). The Protein-RNA complex was eluted from the resin using T4 PNK buffer containing 200 mM imidazole, precipitated with 15% Trichloroacetic acid (TCA) and 2 µg bovine serum albumin (BSA), washed with ice cold acetone, air-dried and resuspended in NuPAGE LDS sample buffer. The sample was heated at 70°C for 10 minutes, electrophoresed on a NuPAGE Novex 4-12% Bis-Tris gel, transferred to nitrocellulose and autoradiographed. A band corresponding approximately to the molecular weight of Utp14 was excised, treated with Protease K (New England Biolabs) for 2 hours at 55°C and the freed RNA was extracted with phenol: chloroform and ethanol precipitated.

Library preparation followed an established protocol for ribosome foot printing (Ingolia et al. 2012). cDNA synthesis was done with the primer (5′-(Phos)-AGATCGGAAGAGCGTCGTGTAGGGAAAGAGTGTAGATCTCGGTGGTCGC-(SpC18)-ACTCA-(SpC18)-<u>TTCAGACGTGTGCTCTTCCGATCT</u>ATTGATGGTGCCTACAG-3’) and either EpiScript RT (Epicentre) or SuperScript III (Invitrogen). Reactions were arrested and RNA hydrolyzed by the addition of NaOH to 100 mM and heating at 98°C for 20 minutes. cDNA was precipitated with ethanol, and resuspended in water. Urea loading buffer (Novex) was added to 1X and the sample was denatured at 80°C for 10 minutes. The cDNA product was electrophoresed on a 10% Novex TBE-Urea gel and extracted in TE, followed by precipitation with isopropanol. The purified cDNA product was circularized using CircLigase (Epicentre) incubated at 60°C for 2 hours, then heat-inactivated at 80°C for 10 minutes. To add adaptors to the first dataset, the circularized product was amplified using Phusion DNA polymerase and oligonucleotides AJO 1986 (5’-AATGATACGGCGACCACCGAGATCTACAC-3’) and ScripMiner Index Primer (#11 for mock and #12 for Utp14-HTP). For the second dataset, the circularized product was initially amplified using Phusion DNA polymerase and flanking oligonucleotides AJO 2299 (5’-TACACGACGCTCTTCCGATC -3’) and AJO 2301 (5’-CAGACGTGTGCTCTTCCGATC -3’). The samples were gel purified as described above, resuspended in water and a subsequent PCR was done to add adaptor sequences using Phusion DNA polymerase and the oligonucleotides AJO 2352 (5’-AATGATACGGCGACCACCGAGATCTACACTCTTTCCCTACACGACGCTCTTCCGATCT -3’) and a ScriptMiner Index Primer (#2 for mock and #4 for Utp14-HTP). The samples were electrophoresed and purified from the gel as described above and resuspended in water.

The resultant cDNA libraries were sequenced on an Illumina MiSeq platform. The single-end reads were processed using fastx_trimmer and fastx_clipper (http://hannonlab.cshl.edu/fastx_toolkit/) to discard low-quality reads and adapter sequences, respectively. The processed reads were aligned to the yeast genome (Ensembl, version R64-1-1) using Bowtie2 (Langmead and Salzberg 2012). The resultant files were analyzed using pyReadCounters and pyPileup (Webb et al. 2014).

### Affinity-purification

Cell growth for affinity-purification are described in the sections below. All steps were carried out at 4°C unless otherwise noted. For mass spectrometry, cells were thawed, washed, and resuspended in one volume of Lysis Buffer (20 mM HEPES-KOH pH 8.0, 110 mM KOAc, 40 mM NaCl, 1mM PMSF and benzamidine, and 1 µM leupeptin and pepstatin). For northern blot analysis, DEPC-treated and nitrocellulose-filtered reagents were used, and cells were resuspended in 1.5 volume of Lysis Buffer. Extracts were prepared using glass beads and clarified by centrifugation at 18,000x*g* for 15 minutes. Clarified extracts were normalized according to A_260_, and TritonX-100 was added to a final concentration of 0.1% (v/v). Normalized extract was incubated for 90 minutes with rabbit IgG (Sigma) coupled to Dynabeads (Invitrogen). The beads were prepared as described (Oeffinger et al. 2007). Following binding, the beads were washed twice in Wash Buffer (Lysis Buffer supplemented with 0.1% TritonX-100) and once with in the Wash Buffer containing 5mM βME at 16°C prior to resuspension in Elution Buffer (Lysis Buffer supplemented with 5 mM βME). For RNA purification, the Elution Buffer was supplemented with 1 U/µL Murine RNase Inhibitor (New England Biolabs). The bound bait-TAP containing complexes were eluted by addition of homemade TEV protease and incubated for 90 minutes at 16°C. The resultant eluates were handed as described in the sections below.

### Northern blot analysis

For Utp14-TAP affinity-purifications, AJY3243 was transformed with the plasmids pAJ4176, pAJ4177, pAJ4178, or pRS415, and YS360 was transformed with pAJ4176. For the Enp1-TAP affinity-purifications, AJY2665, AJY4258, and BY4741 were transformed with pRS416, and AJY4257 was transformed with pRS416, pAJ3422, or pAJ3426. Cell cultures were diluted into in the appropriate SD media containing 2% glucose at a starting OD_600_ of 0.1 and cultured for either 7 hours or grown to mid-exponential phase before collection. Cells were stored at −80°C prior to lysis. Affinity-purifications were performed as described above. Affinity-purified and whole cell extract (WCE) RNAs were isolated using the acid-phenol-chloroform method as described (Zhu et al. 2016). RNAs were separated by electrophoresis through 1.2%-agarose MOPS 6% formaldehyde gel for four hours at 50 volts. Northern blotting was performed as described (Li et al. 2009) using the oligo probes listed in Figure 6 legend, and signal was detected by phosphoimaging on a GE Typhoon FLA9500.

### Mass spectrometry and analysis

For Utp14-TAP affinity-purifications, AJY3243 was transformed with the plasmids pAJ4176, pAJ4177, pAJ4178, or pRS415. Cell cultures were diluted into the appropriate SD media containing 2% glucose at a starting OD_600_ of 0.1 and cultured for either 7 hours or to mid-exponential phase before collection. For the Enp1-TAP affinity-purifications, AJY2665, AJY4257, and AJY4258 cultures were diluted into YPD at a starting OD_600_ of 0.1 and cultured for either 14 hours or to mid-exponential phase before collection. Cells were stored at −80°C prior to lysis. Affinity-purifications were done as described above. To isolate factors associated with only pre-ribosomal particles for mass spectrometry, the eluate was overlaid onto a sucrose cushion (15% D-sucrose, 20 mM HEPES-KOH pH 8.0, 110 mM KOAc, 40 mM NaCl) then centrifuged at 70,000 rpm for 15 minutes on a Beckman Coulter TLA100 rotor.

To perform peptide identification by mass spectrometry, we loaded approximately even amounts of protein from the pellet fraction onto a NuPAGE Novex 4-12% Bis-Tris gel. Proteins were electrophoresed slightly into the gel then stained with Coomassie. A small gel slice containing the proteins was excised and dehydrated with acetonitrile, reduced with 10 mM DTT, then alkylated with 50 mM iodoacetamide. The gel slice was washed with 100 mM ammonium bicarbonate then dehydrated with acetonitrile. In-gel digestion was performed using trypsin (Peirce) in 50 mM ammonium bicarbonate overnight at 37°C. Peptides were extracted with 5% (w/v) formic acid treatment, then with 1:2 (v/v) 5% formic acid : 100% acetonitrile treatment. These solutions were combined with the trypsin digest solution and desalted. The resultant peptides were run for one hour on a Dionex LC and Orbitrap Fusion 1 for LC-MS/MS.

Mass spectrometry data were processed in Scaffold v4.8.3 (Proteome Software, Inc.), and a protein threshold of 99% minimum and 2 peptides minimum, and peptide threshold of 0.1% FDR was applied. The data were exported to Microsoft Excel then custom Python 2.7 scripts were used to calculate the relative spectral abundance factor (RSAF) for each protein by dividing the total number of spectral counts by the molecular weight. For each sample, the RSAF value of each protein was normalized to the mean RSAF value of the UTP-B sub-complex in Microsoft Excel to reflect relative stoichiometry as done previously (Zhang et al. 2016). Supplemental File 2 contains relevant spectral counts and processed data from the mass spectrometry experiments.

## ACKNOWLEDGEMENTS

We wish to thank J. Recchia-Rife for his assistance with cloning and J.A. Hussmann for his initial assistance with the CRAC data analysis. We thank Dr. E. Petfalski for strain YS360

## DATA AVAILABILITY

All relevant sequencing data have been deposited in the Gene Expression Omnibus (GEO) database (http://www.ncbi.nlm.nih.gov/geo/) with the accession number XXXXX. Python scripts are available upon request.

